# Long-read RNA-seq demarcates *cis*- and *trans*-directed alternative RNA splicing

**DOI:** 10.1101/2024.06.14.599101

**Authors:** Giovanni Quinones-Valdez, Kofi Amoah, Xinshu Xiao

## Abstract

Genetic regulation of alternative splicing constitutes an important link between genetic variation and disease. Nonetheless, RNA splicing is regulated by both *cis*-acting elements and *trans*-acting splicing factors. Determining splicing events that are directed primarily by the *cis*- or *trans*-acting mechanisms will greatly inform our understanding of the genetic basis of disease. Here, we show that long-read RNA-seq, combined with our new method isoLASER, enables a clear segregation of *cis*- and *trans*-directed splicing events for individual samples. The genetic linkage of splicing is largely individual-specific, in stark contrast to the tissue-specific pattern of splicing profiles. Analysis of long-read RNA-seq data from human and mouse revealed thousands of *cis*-directed splicing events susceptible to genetic regulation. We highlight such events in the HLA genes whose analysis was challenging with short-read data. We also highlight novel *cis*-directed splicing events in Alzheimer’s disease-relevant genes such as *MAPT* and *BIN1*. Together, the clear demarcation of *cis*- and *trans*-directed splicing paves ways for future studies of the genetic basis of disease.

## Main

Alternative splicing is an essential mechanism in eukaryotic cells that enables substantial transcriptomic diversity, playing critical roles in biology and disease^1,2^. Splicing is a closely regulated process, primarily governed by the interactions between *cis*-regulatory elements and *trans*-acting factors, such as RNA binding proteins (RBPs). Disruption of *cis*-regulation of splicing by genetic variants is a primary link between genotypes and disease^2,3^. On the other hand, *trans*-factors are essential splicing regulators, orchestrating the substantial diversity of splicing profiles across cell types, tissues, and developmental stages^1,4,5^. In addition, the two aspects of splicing regulation are closely intertwined since interactions between the *cis*- and *trans*-regulators are essential for their function.

In recent years, an increasing amount of effort has been dedicated to studying the association between genetic variants and splicing. Using methods such as large-scale functional screens^6–9^, splicing quantitative trait loci (sQTL)^10^, allele-specific splicing^11^, machine learning^12–15^ or large-scale data-driven approaches^16,17^, many genetic variants have been uncovered that are associated with or cause splicing alterations. Such molecular associations can be further exploited to prioritize candidate causal genes for diseases or traits following genome-wide association studies (GWAS)^18,19^. Based on these studies, splicing is emerging as an essential molecular trait that informs the genotype-phenotype relationships.

Interestingly, recent studies showed that not all splicing events are equally susceptible to aberrant disruption by genetic variants^20,21^. Certain exons, named hotspot exons, were shown to be prone to exon skipping and enriched with splice-disrupting variants^21^. The vulnerability of an exon to splice-altering variants may depend on its sequence context and basal exon inclusion level^20,22^. For exons genetically regulated by *cis*-variants, many exhibit signs of positive selection^6,23^ with enrichment in specific biological processes, such as immune-related pathways^11,23^. Similarly, species-specific splicing events are more often *cis*-directed than *trans*-directed^24–26^. Together, these studies suggest that certain exons are more prone to *cis*-disruption than others, although every exon is controlled by both *cis*- and *trans*-acting regulators.

Here, we demonstrate the efficacy of long-read RNA-seq data in discerning *cis*- and *trans*-directed splicing events. Specifically, the *cis*-directed events are characterized by allele-specific alternative splicing patterns, which can be readily identified with our new method, isoLASER. In contrast, *trans*-directed events exhibit no linkage between haplotypes and splicing. We map the global landscape of *cis*-and *trans*-directed splicing by analyzing long-read RNA-seq data from human and mouse samples. Notably, our approach successfully uncovers *cis*-directed splicing in the highly polymorphic HLA system, which is difficult to achieve with short-read sequencing data. Applied to data derived from Alzheimer’s patients, we report disease-specific *cis*-directed events, including those in the HLA family and other genes known to contribute to the pathogenesis of the disease. This study delineates *cis*- and *trans*-directed splicing events, opening avenues for exploring disease mechanisms and potential therapeutic targets.

## Results

### Long-read RNA-seq uncovers cis- and trans-directed splicing events

Long-read RNA-seq possesses a unique strength in uncovering full-length isoforms of each gene and, when combined with genotype information, may unveil haplotype-specific splicing and other alternative RNA processing events. To explore this application, we examined long-read RNA-seq data generated by the ENCODE consortium in K562 cells using the PacBio Sequel II platform^27^. As an example, for the gene *RIPK2*, the long reads can be separated into two haplotypes (H1 and H2) based on the genotype of a heterozygous SNP (Fig. 1a). Strikingly, this analysis clearly uncovered two classes of alternatively spliced (AS) events. For one class of events (highlighted in blue, Fig. 1a), exon inclusion was observed almost exclusively in only one haplotype (H1). In contrast, the second class of AS events (highlighted in red, Fig. 1a) had approximately the same level (56% in H1 vs. 61% in H2) of exon inclusion in the two haplotypes. Thus, the haplotype-specific nature of the first class of exons reflects a dominating role of *cis*-regulatory genetic variants on splicing (with the causal variants being unknown), hereby referred to as “*cis*-directed” exons. For the second class of events, genetic modulation is not the dominant factor in the splicing regulation. Hence, we call this class “*trans*-directed” exons. Note that this classification reflects the prevailing regulatory mechanisms of an exon in a specific cellular context, which does not override the fact that a combination of *cis*-elements and *trans*-acting factors controls the splicing event.

**Figure 1.**
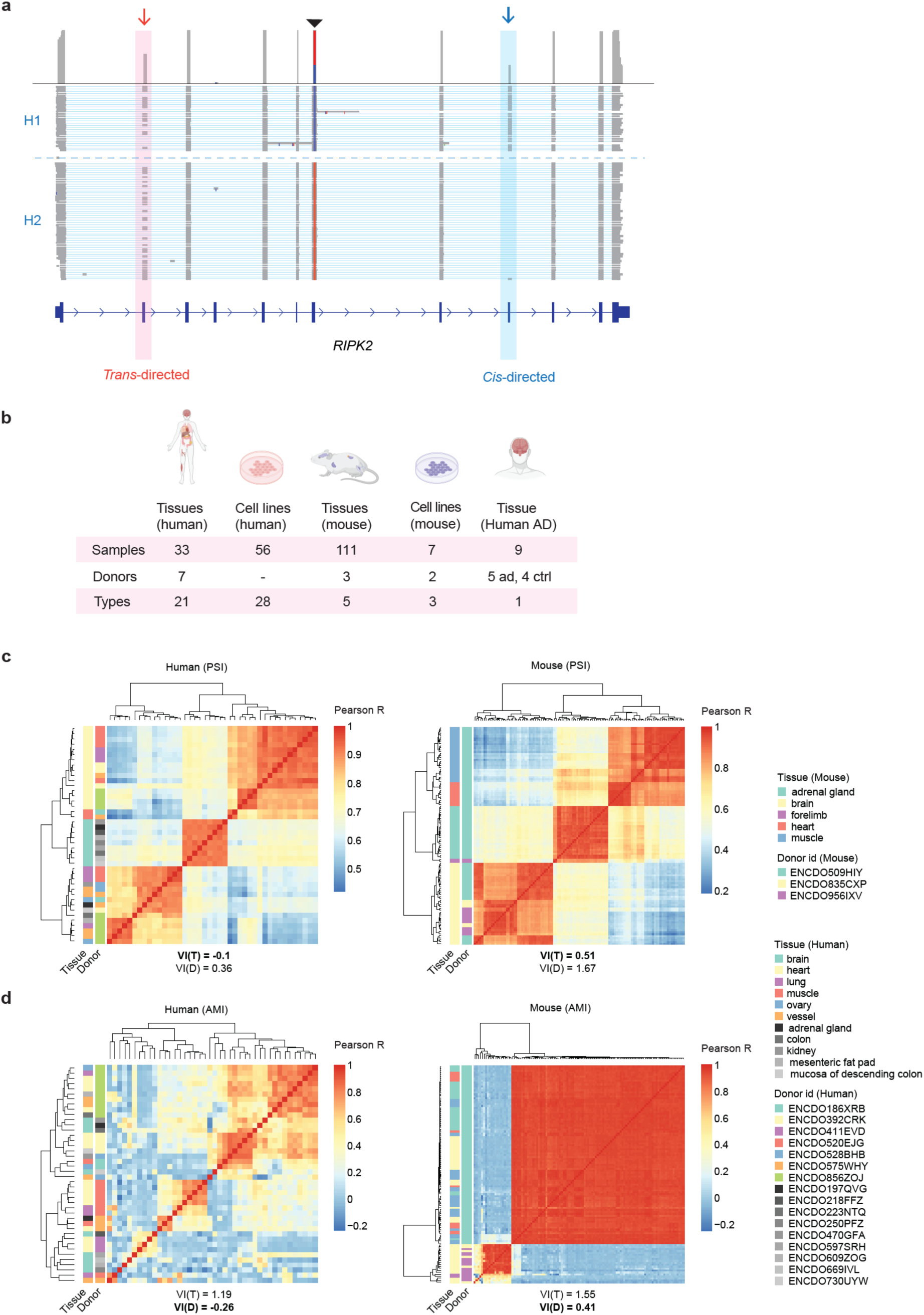
isoLASER and dataset overview. **a,** Examples of a *cis-* and a *trans*-directed exonic part in the gene *RIPK2* in K562 cells. Reads were grouped by haplotype (H1 vs. H2) based on the genotype of an exonic SNP (black arrow head). The *cis*- and *trans*-directed events are highlighted in blue and pink, respectively. **b,** Overview of ENCODE4 long-read RNA-seq datasets used in this study. For each category, the numbers of samples, donors, and tissue or cell types are shown. ad: Alzheimer’s disease; ctrl: controls. **c,** Heatmap of pairwise Pearson’s correlations using the PSI of all alternatively spliced exonic parts in human (left) and mouse (right) tissues. Tissue types and donor IDs are shown. Donors that only contributed one tissue are colored in shades of gray. Similarly, tissues present in only one donor are colored in shades of gray. Meila’s variation of information (VI) is given for the clustering consistency with the tissue VI(T) and donor VI(D). Lower values indicate a higher correlation. The lowest VI value is bolded. **d,** Similar to **c,** but using the adjusted mutual information (AMI) for Pearson’s correlation.

Next, we analyzed all long-read RNA-seq data from human and mouse tissues/cell lines generated by the ENCODE consortium^27^ (Fig. 1b). As an initial overview of global splicing profiles, we calculated percent-spliced-in (PSI) values of AS exons and clustered the human or mouse tissues based on the Spearman correlation of their PSI values. Consistent with previous literature, the samples were clustered primarily according to their tissue of origin^28^ (Fig. 1c). Next, we calculated a metric, adjusted mutual information (AMI), to quantify the allelic linkage between genetic variants and splicing levels of alternative exons (Methods). Remarkably, the AMI-based clustering segregated the samples primarily based on donor identity rather than tissue of origin (Fig. 1d). This analysis included all AS exons that resided in reads harboring heterozygous SNPs, not restricted to exons with evident haplotype-specific splicing as highlighted in Fig. 1a. This result strongly suggests that an individual’s genetic background plays an important role in shaping their overall splicing profile. Additionally, this observation underscores the potential widespread distinction of *cis*- and *trans*-directed splicing events, as illustrated in Fig. 1a.

### The isoLASER method to detect cis- and trans-directed events in long-read RNA-seq

Next, we aimed to globally identify *cis*- and *trans*-directed events by developing a new method called isoLASER (isoform-Level analysis of Allele-Specific processing of Exonic Regions). Although the final goal is to analyze allelic linkage of splicing, isoLASER provides a one-stop solution by performing three major tasks: *de novo* variant calls, gene-level phasing of variants, and linkage testing between the phased haplotypes and the alternatively spliced exonic segments (i.e., allelic linkage of splicing; Fig. 2a, Methods).

**Figure 2.**
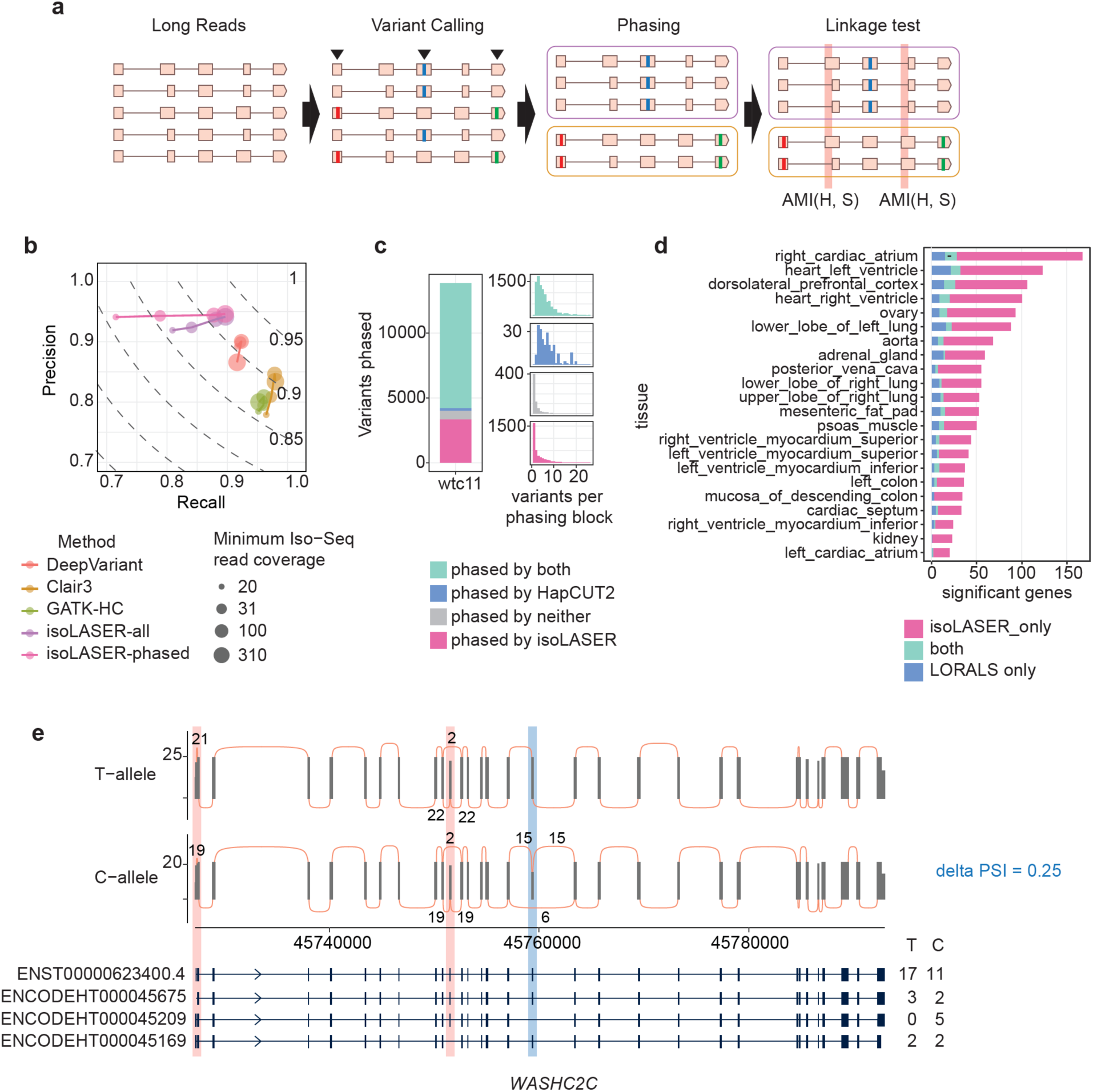
isoLASER overview and benchmarking. **a,** Illustration of the three major steps in isoLASER: de novo variant calling, haplotype phasing and linkage analysis. **b,** Variant calling performance of isoLASER, GATK HaplotypeCaller (HC), DeepVariant and Clair3 with WTC11 data. Precision and Recall are shown for different read coverage cutoffs. The dashed curves indicate F1 score thresholds. Variant calling performance is shown for all variants identified by isoLASER (all) or for the variants used for phasing (phased). **c,** Heterozygous variant phasing by isoLASER and HapCUT2. The total numbers of variants phased by either or both methods are shown. The number of variants contained in each phasing block is also shown. WTC11 data were used in this analysis. **d,** Number of genes with a *cis*-directed splicing event identified by isoLASER or with allele-specific transcript expression identified by LORALS in the ENCODE human tissue dataset. **e,** A *cis*-directed event identified in the gene *WASHC2C.* No allele-specific transcript expression was identified for this gene using LORALS. The *cis*- and *trans*-directed exonic parts are highlighted in blue and red, respectively. The delta PSI of the *cis-*directed event calculated by isoLASER is shown on the side.

First, isoLASER conducts variant calling using the long-read RNA-seq data. It uses a local reassembly approach based on de Bruijn graphs to identify nucleotide variation at the read level, followed by a multi-layer perceptron classifier to discard false positives. This classifier achieves a training performance with the AUC between 0.92 and 0.99 for the receiver operating characteristic (ROC) curve and between 0.86 and 0.99 for the precision-recall curve (Fig. S1a)^29^. To evaluate its performance on testing data, we followed the variant calling benchmark protocol in de Souza et al^30^ (Methods). Using genotyped long-read RNA-seq from WTC11 cells^30^, isoLASER achieved similar F1 scores as GATK’s Haplotype Caller^31^, DeepVariant^32^, and Clair3^33^ (with their corresponding pre-processing) but with superior precision (Fig. 2b), which is a desirable feature in typical applications.

Following variant calling, isoLASER next carries out gene-level phasing to identify haplotypes. Briefly, after read-level variant calls, an approach based on k-means read clustering is employed, using the variant alleles as values and weighted by the variant quality score (Methods). This step simultaneously phases the variants and groups individual reads into their corresponding haplotypes. We compared the phasing performance of isoLASER with that of HapCUT2^34^, a method designed to precisely phase variants using different sequencing protocols including single-molecule long-read sequencing. Using the same variant calls and alignment files from WTC11 cells, we compared the phasing rate and consistency between the two methods. Over 99% of genes containing multiple heterozygous variants phased by HapCUT2 were consistently phased by isoLASER (Fig. 2c), supporting the effectiveness of isoLASER in this step. Additionally, isoLASER phases haplotype blocks that contain few variants more frequently than HapCUT2.

Subsequently, the allelic linkage of splicing is analyzed for exonic segments (i.e., exonic parts) that are non-overlapping, unique exonic regions with distinct splicing patterns from each other. Exonic parts represent the basic units of exons that reflect local alternative splicing events, which enables event-specific genetic association analysis. For each gene, the allelic linkage between phased haplotypes and exonic parts was quantified by the AMI (Methods). We simulated unlinked events for linkage testing to determine the AMI cutoffs at different read coverage levels to control false positives (Fig. S1b, Methods). Based on this analysis, we defined *cis*-directed events as those with AMI greater than 99% of the simulated background and an absolute delta PSI (PSI difference between haplotypes) greater than 5%. In contrast, *trans*-directed events as those with AMI smaller than 95% at the corresponding read coverage level. Other events are defined as ambiguous. A quantile-quantile plot shows that *cis*-directed events identified in our dataset had a much higher AMI than expected by chance (Methods) (Fig. S1c), likely reflecting the stringency in calling *cis*-directed events.

We compared allelic linkage results of isoLASER to those of LORALS, a recently published method for allele-specific analysis of long-read RNA-seq data^35^. We applied both methods to ENCODE data generated from human tissues. isoLASER identified substantially more genes with allele-specific splicing events (i.e., *cis*-directed events) than LORALS (Fig. 2d). Unlike isoLASER, LORALs requires previously genotyped and phased data. A major conceptual distinction between the two methods is that LORALS focuses on isoform-level allelic linkage whereas isoLASER tackles individual exonic parts. Since human genes often harbor multiple AS events, whose combinatorial inclusion may result in a large number of isoforms, it is expected that focusing on local AS events affords greater sensitivity than isoform-level approaches. As an example, the gene *WASHC2C* has one exonic part spliced in an allele-specific manner identified by isoLASER (blue highlighted, Fig. 2e). In contrast, none of the 4 annotated transcripts showed significant allele-specific bias that was identifiable by LORALS. Furthermore, isoLASER quantifies the splicing difference between haplotypes using the delta PSI, providing a more interpretable metric of the splicing difference between haplotypes.

In the above description, isoLASER was applied to each sample individually. Applicability to a single sample is a major advantage of allele-specific analysis. Nonetheless, leveraging the existence of multiple samples, if available, is also important. We next extended isoLASER to jointly analyze multiple samples to achieve two objectives: (1) to prioritize likely functional variants and (2) to identify *cis*-events that were otherwise untestable or weak in individual samples. This approach allowed us to leverage the diverse genetic backgrounds of various donors (Methods, Fig. S1d). Henceforth, we will refer to this joint analysis as isoLASER-joint and single-sample analysis as isoLASER.

### Cis-directed events are enriched in immune-related genes

We next applied isoLASER to all samples in Fig. 1b. In each tissue sample or cell line (human or mouse), we identified 2 to 946 *cis*-directed events (Fig. 3a, S2a-c, Table S1). Across human (mouse) tissues, a total of 2,047 (4,679) unique *cis*-directed exonic parts in 1,203 (2,341) unique genes were identified, and for human (mouse) cell lines, a total of 3,312 (1,341) unique exonic parts and 1,703 (778) unique genes were detected. About 13% of the *cis*-directed exonic parts were not annotated in GENCODE, thus denoting novel exonic parts. As expected, samples with higher coverage had more *cis*-directed splicing events (Fig. 3a, S2a-c). Moreover, samples from F1 hybrid mouse strains demonstrated higher levels of *cis*-directed events than F0 inbred samples (Fig. S2b,c). This observation aligns well with the number of heterozygous variants in hybrid mice, facilitating the detection of *cis*-regulated splicing.

**Figure 3.**
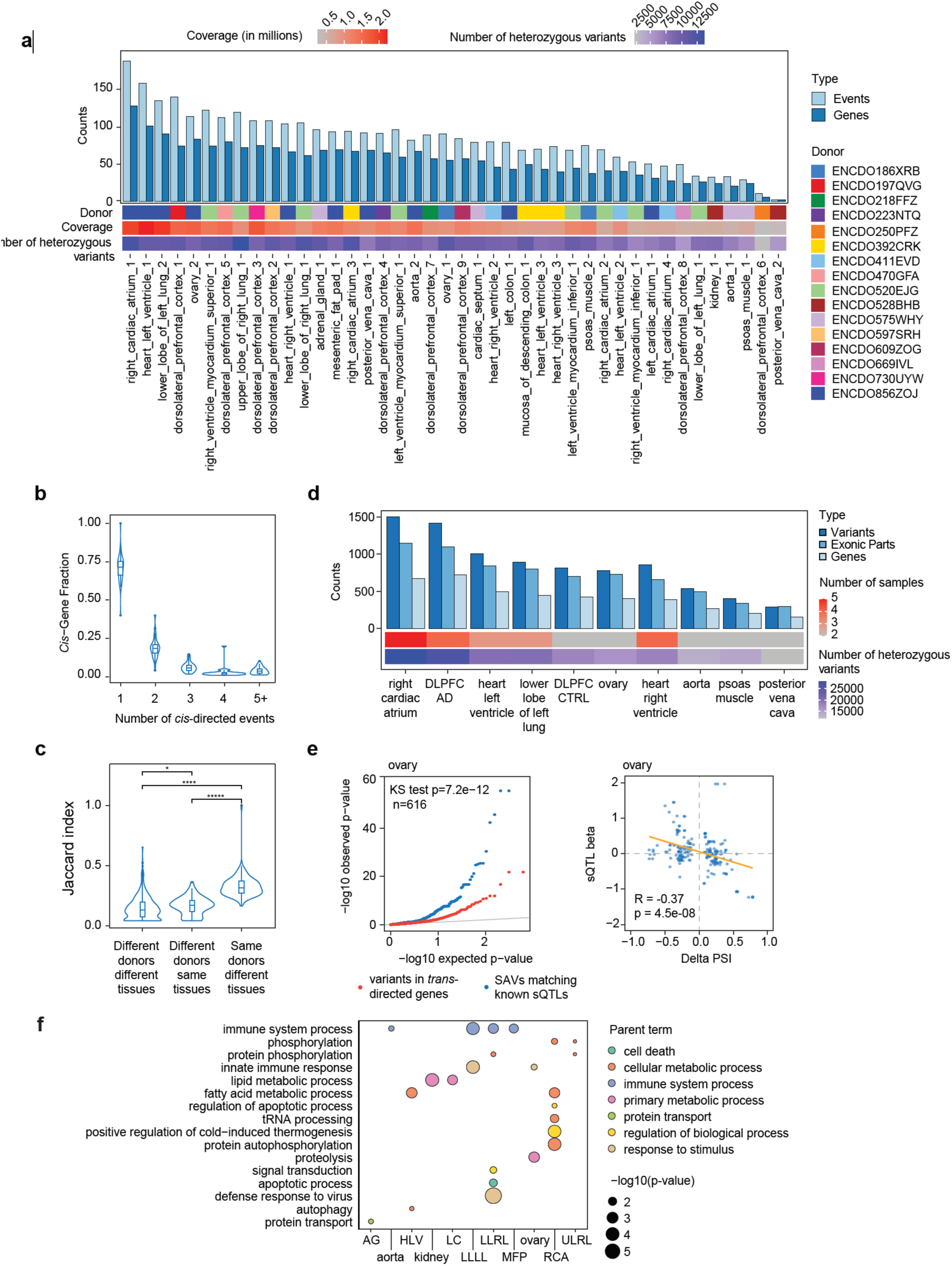
Frequency of *cis*-directed events in human tissue samples. **a,** Number of unique *cis*-directed exonic events and genes detected in each human tissue sample. The number of heterozygous variants, total read coverage, and donor membership are shown for each sample. **b,** Fraction of genes with N (1, 2, 3, 4, 5+) *cis*-directed exons among all genes with any *cis*-directed events. This fraction was calculated for each sample and visualized for all human tissue samples. **c,** Sharing of *cis*-directed events, measured by the Jaccard index, between samples originating from the same or different individuals. **d,** Number of variants, *cis*-directed exonic parts, and genes detected by merging human samples of the same tissue type using isoLASER-joint. DLPFC: dorsolateral prefrontal cortex. **e,** Left: quantile-quantile plot showing the distribution of GTEx sQTL p-values in ovary. Blue: splicing-associated variants (SAVs) discovered by isoLASER-joint that had sQTL p values (regardless of significance) in similar GTEx tissues. Red: variants in *trans*-directed genes that had sQTL p values. Right: Correlation between delta PSI of the variants calculated by isoLASER and the genotype beta parameter from the linear regression calculated in GTEx sQTL mapping. Intron usage instead of PSI was used by GTEx as the splicing phenotype. Thus, a negative correlation is expected. The p-value and regression coefficient were calculated using Pearson’s correlation. The trendline was fit using a linear regression model. **f,** Gene Ontology (GO) enrichment analysis of genes containing *cis*-directed events from human tissues. Samples derived from the same tissue type were combined. GO terms are color-coded based on their parent terms. Tissue names: AG = adrenal gland; HLV = Heart Left Ventricle; LC = left colon; LLLL = lower lobe of the left lung; LLRL = lower lobe of the right lung; MFC = mesenteric fat pad; RCA = right cardiac atrium; ULRL = upper lobe of the right lung. The terms are sorted by the frequency of significance across tissues; terms enriched in more tissues are positioned higher in the list.

Among genes with *cis*-directed events, 30-40% had two or more such events (Fig. 3b, S2d-f). Notably, *cis*-directed events are more frequently shared among different tissues originating from the same donor compared to those (same or different tissue types) from different donors (Fig. 3c, Fig. S2g-h). This observation is again consistent with the nature of *cis*-directed events where splicing is closely dependent on the genetic background of the sample. It also aligns with the earlier observation that the samples were largely segregated by donors when clustered by AMI (Fig. 1d). In general, *trans*-directed events were more abundant than *cis*-directed events at the gene level (Fig.

S3). Nonetheless, a *trans*-directed event in one sample could be a *cis*-directed event in another (Fig. S3), depending on the genetic background. If all human tissues are considered together, 15% of the testable events were *cis*-directed in at least one tissue, with 68% being *trans*-directed (the rest being ambiguous).

We next applied isoLASER-joint to each tissue/cell type (requiring ≥2 samples). The number of *cis*-directed events increased compared to the single-sample analysis (Fig. 3d, S4a,b), as expected. In addition, the genetic variants associated with *cis*-directed events (Table S2) had significantly lower p-values in the sQTL analysis reported by the GTEx study^10^, compared to the p-values of matched controls (Fig. 3e left; Fig. S4c). Furthermore, the effect sizes (delta PSI from isoLASER) of the genetic variants overlapping sQTLs were significantly correlated with the genotype beta parameter from the sQTL regression (Fig. 3e right, Fig. S4d). These results indicate that isoLASER-joint achieves an sQTL-type of analysis, requiring only limited sample sizes.

Next, we conducted a Gene Ontology (GO) analysis of genes containing *cis*-directed splicing events in at least one human tissue sample (Methods). The analysis revealed significant terms, including immune system process and innate immune response (Fig. 3f). Among immune-related genes are *PKR, CD44, LILRB3, YPEL5,* members of the immunoglobulin superfamily (*CD146, BTN3A2*) and members of the human leukocyte antigen (HLA) family (Table S3). The enrichment of *cis*-directed events in immune-related genes supports the idea that genetically regulated splicing is an important mechanism underlying the genetic architecture of immune diversity^36^.

### isoLASER uncovers allele-specific splicing of HLA genes and enables HLA typing

Among the above immune-related genes that harbored *cis*-directed events are multiple members of the HLA family, such as *HLA-A, HLA-DPB1, and HLA-DQA1* (Fig. 4a, Table S4). HLA molecules present peptides to immune cells, thus playing a pivotal role in T-cell activation and effective immune response^37,38^. As the most polymorphic genetic system in humans, HLA genes contribute significantly to heritability, surpassing all other known loci combined^39,40^. Although HLA typing is essential for understanding the immune response in health and disease, the roles of alternative splicing in HLA expression remain poorly understood.

**Figure 4.**
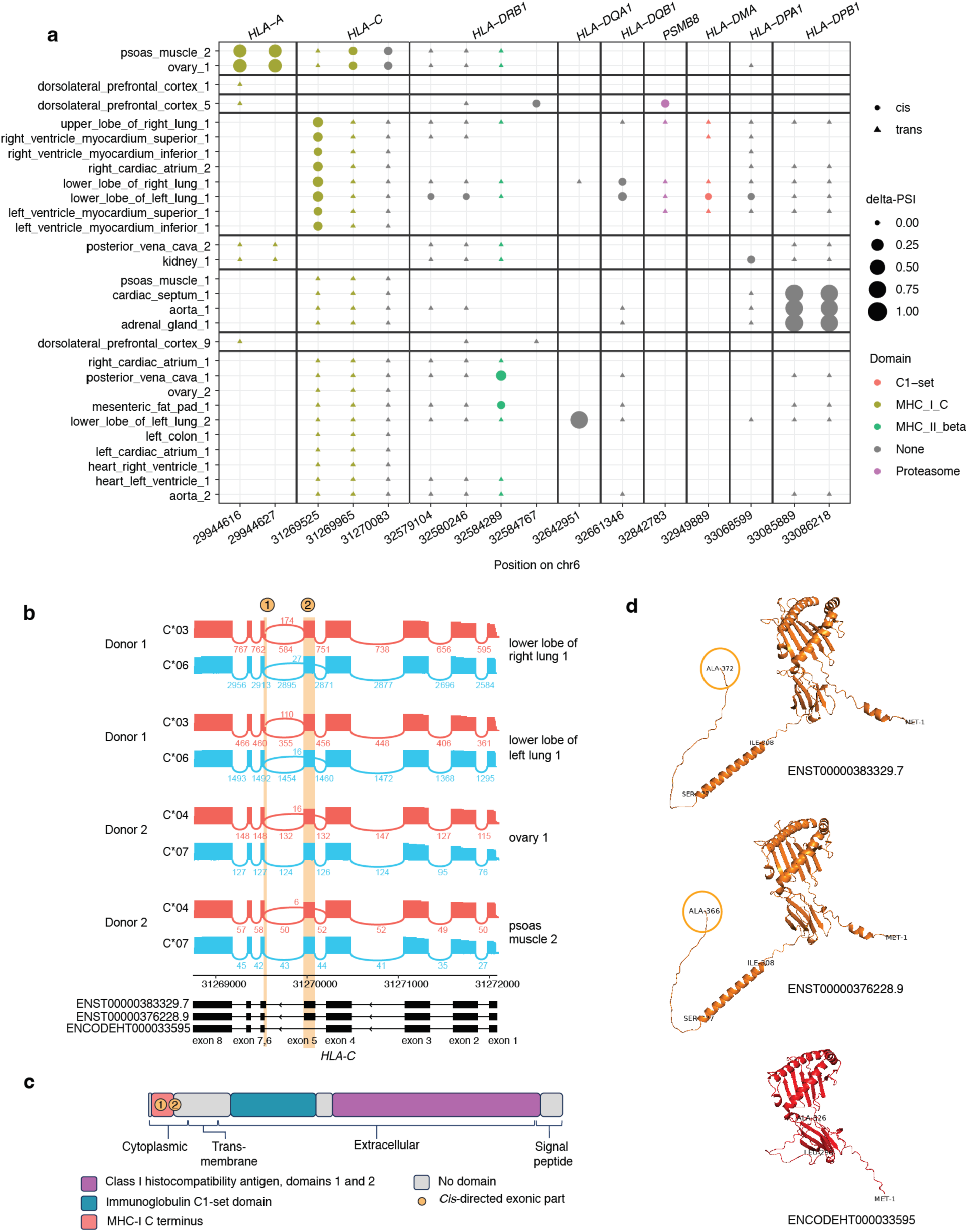
Allele-specific splicing of HLA genes. **a,** Summary of *cis*-directed events in the HLA gene family. The coordinates on the x-axis indicate the starting position of the exonic parts and are grouped by gene. The samples on the y-axis are grouped by donor. The size of the dot indicates the effect size (delta PSI) of the events. Both *cis*- and *trans*-directed events are shown (*trans*-directed events have close to 0 delta PSI). The dots are colored to denote the Pfam protein domains overlapping the *cis*-directed events. **b,** The gene *HLA-C* contains two *cis*-directed events, one specific to donor 1 (event labeled as1) and two specific to donor 2(event labeled as 2). These donor-specific events are consistent across different tissues of the same donor. The HLA-type corresponding to the major allele group is shown to the side of each haplotype. **c,** Protein domain (Pfam) and cellular localization diagram of the *HLA-C* gene with the relative position of the *cis*-directed exonic parts. **d,** AlphaFold predictions of three selected isoforms that contain or lack the *cis*-directed exonic regions. The first and last amino acids of the entire protein and of the *trans*-membrane domain were labeled as references. The position of the last amino acid (circled) indicates the length difference between the first two isoforms (6 amino acids difference).

IsoLASER allows HLA typing and a detailed analysis of allele-specific splicing patterns of each HLA gene. We observed individual-specific *cis*-directed events across multiple HLA genes, with many of them overlapping annotated protein domains from the Pfam database^41^ (Fig. 4a). For example, *HLA-C* contains two individual-specific *cis*-directed events consistently observed across tissues of each individual (Fig. 4b). To gain a more in-depth view of this gene, we matched the haplotypes identified by isoLASER in human tissues with known HLA-C alleles from the IPD-IMGT database^42^. This analysis revealed consistent splicing patterns for each HLA-C allele irrespective of the tissue of origin (Fig. S5). Specifically, the C*04 and C*06 alleles consistently exhibited some degree of exon 5 skipping. The C*03 allele had an alternative splice site in exon 6, with a relative usage of approximately 25%, whereas the C*07 allele exhibited no alternative splicing. The *cis*-directed events in *HLA-C* may have important functional implications. Specifically, they overlap with the MHC-I C-terminus domain situated in the cytoplasm (exon 5), as well as a transmembrane alpha helix (Fig. 4c). AlphaFold predictions of isoforms with or without the *cis*-directed events revealed substantial alterations to the resulting protein products (Fig. 4d). Specifically, event 1, yielding isoform ENST00000383329.7 through the usage of an alternative 3’ splice site on exon 6, results in the elongation of the MHC-1 C terminus by 6 amino acids, compared to ENST00000376228.9. Event 2, yielding isoform ENCODEHT000033595 through the exclusion of exon 5, results in the deletion of a large section of the transmembrane helix causing complete loss of the helix structure.

Other genes in the HLA family also contain *cis*-directed events in functional domains, such as *HLA-A*, *HLA-DMA* and *HLA-DPB1* (Fig. 4a, Fig. S6a-i). Similar to *HLA-C*, the haplotypes uncovered in the reads of the other genes also matched with known HLA alleles. Thus, the long-read RNA-seq data generally facilitate HLA typing in the RNA. *Cis*-events in these genes may also have important functional implications, such as by affecting the C-terminus of the HLA-A protein (Fig. S6a-c), the C1-set domain of HLA-DMA that is in charge of T-cell recognition^43^ (Fig. S6d, e, f), or the extension/shortening of the signal peptide in HLA-DPB1 (Fig. S6g,h, i). Altogether, isoLASER helps to uncover a number of allele-specific splicing events in HLA genes, with potential implications for protein function, which complements the traditional methods of DNA-centric HLA typing.

### Allele-specific splicing of HLA genes in AD patients

Recently, HLA genes are increasingly recognized as important contributors to Alzheimer’s disease (AD)^44^. Using the data generated from the dorsolateral prefrontal cortex (DLPFC) tissue of 5 AD patients and 4 controls (Fig. 1b), we asked whether allele-specific splicing events in HLA genes were observed in AD and whether such events were specific to AD compared to controls. Combining the results of isoLASER and isoLASER-joint, two of the HLA genes demonstrated allele-specific splicing in the AD samples, including *HLA-C* and *HLA-DMA* (Table S5). To expand this analysis, we also examined another long-read RNA-seq dataset (Oxford Nanopore) generated from the DLPFC of 6 AD patients and 6 controls^45^. This dataset yielded seven HLA genes with allele-specific splicing in the AD samples (Table S5), including the two genes from the ENCODE data (Fig. 5a).

**Figure 5.**
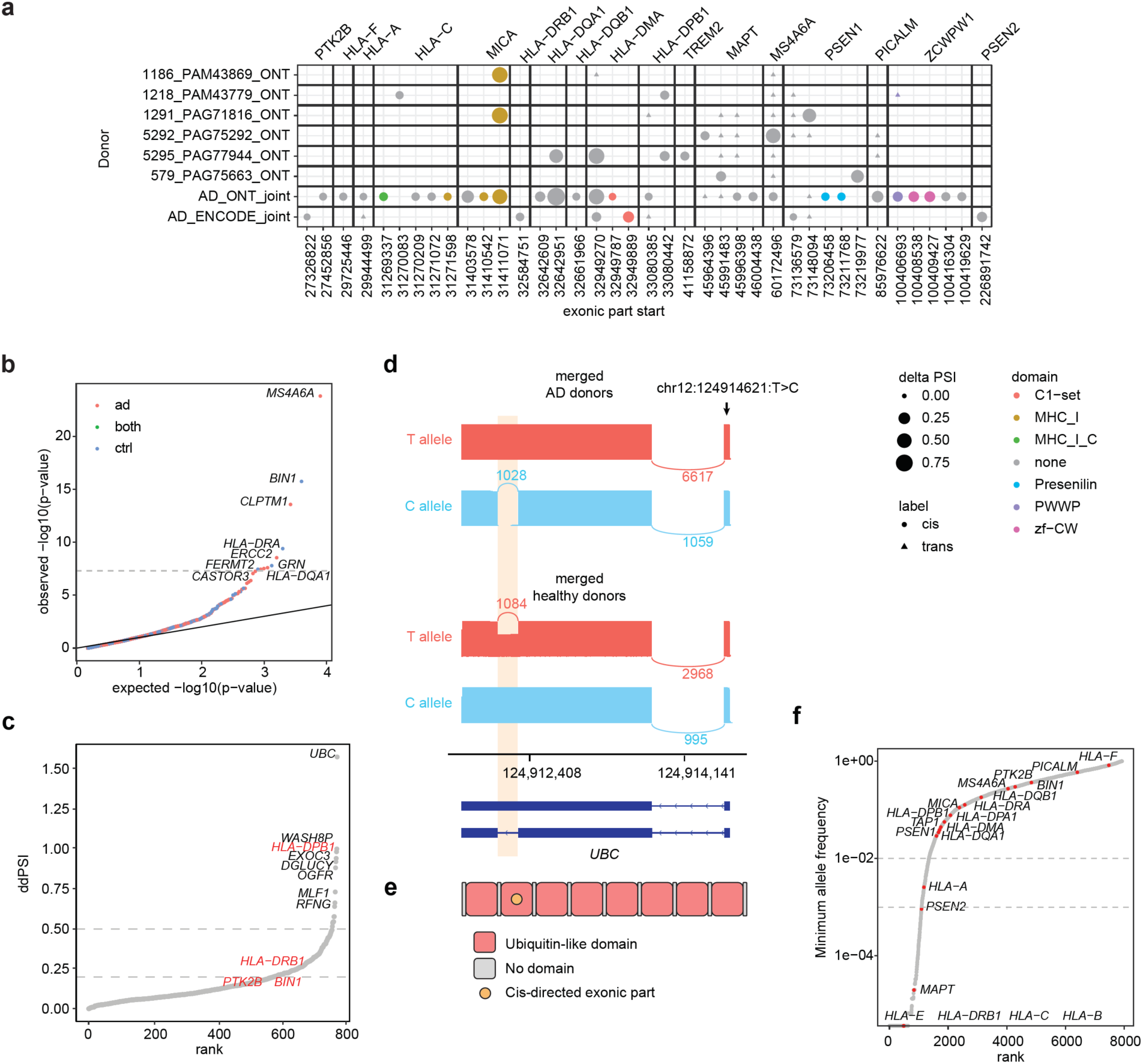
Allele-specific splicing in Alzheimer’s disease. **a,** Summary of *cis*-directed events in the HLA gene family and known AD risk genes in samples from the frontal cortex (Nanopore data) and dorsolateral prefrontal cortex (PacBio data). Events from the sample-level and joint analysis are shown. X-axis are labeled with the coordinates of the starting positions of the *cis*-directed events. **b,** Quantile-quantile plot of the Alzheimer’s GWAS p-values of SAVs identified by isoLASER. The GWAS threshold for significance is shown as a dashed line (5×10^-8^). SAVs identified only in AD, only in controls or in both groups are colored differently. **c,** *Cis*-directed events ranked by their ddPSI values. Gene names are shown for those with a ddPSI value greater than 0.5 or 0.2 (only if the gene was previously identified as an AD risk gene, red). **d,** Differential allele-specific splicing pattern is present in the Ubiquitin C gene (*UBC*), which has the highest ddPSI value shown in **c**. The inclusion of the highlighted intronic region is strongly linked to the variant at position 124914621 on chromosome 12, but the specific linkage pattern is reversed in the two conditions. **e,** Diagram illustrating the repeating ubiquitin-like domains and the *cis*-directed exonic part in the *UBC* gene. **f,** Minor allele frequency (from gnomAD) of the SAVs identified by isoLASER in the AD and control samples. Variants not found in gnomAD were assigned an MAF = 0. Indels not found in gnomAD were discarded. SAVs located in HLA and AD risk genes are labeled in red.

Given the presence of *cis*-directed events in HLA genes, we next asked if other AD-relevant genes also harbored *cis*-directed events. Genes such as *MAPT, PSEN1, MS4A6A, PICALM,* and *ZCWPW1* were found to have such events in at least one sample (Fig. 5a, Fig. S7). Through the isoLASER-joint analysis, we identified the variants associated with these events (splicing-associated variants, SAVs) and overlapped them with the GWAS summary statistics for AD. SAVs located in the genes *MS4A6A, BIN1, HLA-DRA,* and *HLA-DQA1,* among others, passed the GWAS p-value cutoff for significance, suggesting their potential contribution to the disease (Fig. 5b). Although some events were only observed in control donors, the GWAS significance of the SAVs indicates that the risk allele is associated with AD. It is also notable that these variants were not previously found to be associated with splicing through splicing QTL studies^46^ for AD, underscoring the advantage of our method.

We next identified differential allele-specific splicing events between AD and controls in the HLA and AD risk genes in the above two datasets, respectively (Methods). Briefly, we compared the allele-specific effect size (delta PSI) of each SAV and then quantified the difference between AD and controls as the delta-delta PSI (ddPSI). As shown in Fig. 5c, a number of genes showed *cis*-events with significant ddPSI values indicating the potential existence of disease-specific genetic regulation. Multiple genes involved in neurological conditions, such as *UBC,* contain multiple events with high ddPSI values. An intron (highlighted in Fig. 5d) of the *UBC* gene showed 100% retention in the haplotype carrying the T allele of the highlighted variant in AD samples but nearly 0% retention associated with the C allele. Remarkably, this pattern was reversed in control samples, with the C allele associated with 100% intron retention but the T allele with ∼50% retention. The coding region of *UBC* comprises nine tandemly repeated moieties of the ubiquitin domain^47^. The highlighted alternative splicing event removes one ubiquitin domain from the final transcript (Fig. 5e), which can potentially disturb the tightly regulated cellular-free ubiquitin pool equilibrium^48^. Accumulation of misfolded proteins is a hallmark of multiple neurological diseases, including AD. The limited number of free ubiquitins may impair the proper tagging and later degradation of Tau and Amyloid beta aggregates^49,50^.

Lastly, using the gnomAD database^51^ we extracted the minor allele frequency (MAF) of the SAVs identified in this analysis. Over 1,201 variants (15% of the total) were classified as either rare (MAF < 10^-2^) or ultra-rare (MAF < 10^-3^) (Fig. 5f). Multiple SAVs identified in the HLA genes, *MAPT*, and *PSEN2* genes fall under this category. Rare variant analysis is very challenging via genome-wide association methods, such as splicing QTL, due to their low prevalence. Here, our data show that isoLASER-joint can evaluate these variants directly, stemming from the allele-specific nature of this method.

## Discussion

RNA splicing is primarily determined by the contribution of *cis*-acting sequence elements and *trans*-acting splicing factors. Genetic variants may disrupt *cis*-regulatory motifs and alter splicing, constituting an essential link between genotypes and phenotypes. Here, we leverage long-read RNA-seq to demarcate splicing events that are primarily *cis*- or *trans*-directed, using a new method, isoLASER, that performs *de novo* variant calling, gene-level phasing, and allele-specific splicing analysis for long reads.

isoLASER focuses on exonic parts instead of an isoform-level analysis^35,52^. This approach affords a granular view of individual splicing events and the associated isoform diversity. For example, multiple *cis*-directed events may be present in a specific gene, with different levels of allele-specific association. Since many transcript isoforms may be needed to represent the combinatorial inclusion or exclusion of these events, isoform-level analyses may be underpowered and yield insignificant observations. Thus, isoLASER complements previous studies of allele-specific isoform expression^35,52,53^ from the unique perspective of exonic parts. Furthermore, the usage of the adjusted mutual information and delta PSI offers a more intuitive understanding of partial associations compared to other statistical tests. Additionally, isoLASER leverages the read-level associations representing direct evidence of genetically modulated alternative splicing. This principle allows the detection of splicing-associated variants in small cohorts, presenting a viable alternative to splicing QTL mapping. Similarly, it also allows the examination of rare variants, as the only technical requirement is that such variants have sufficient read coverage in a sample. This aspect represents a remarkable advantage over association studies that require modeling or imputation of their effect size and significance.

We demonstrated that the genetic linkage profiles of alternative splicing events are highly individual-specific and maintained across different tissues. Gene Ontology analysis revealed that genes harboring *cis*-directed events are enriched with immune-related functions. This finding is consistent with reports of highly individual-specific immune adaptation through transcript alterations^54–56^. Among genes with critical immune relevance is the HLA gene system, which has been associated with more complex diseases than any other genomic loci^39,40,57^. Due to their high polymorphic nature, allele-typing and allele-specific splicing of HLA genes are challenging to resolve using short reads. Long-read RNA-seq combined with isoLASER allowed us to uncover *cis*-directed splicing events associated with the different HLA allele groups, with some events causing significant disruption to the protein structure. Despite their potential functional impact, allele-specific splicing events in HLA genes were poorly characterized. Our method systematically identifies splicing events associated with the major HLA allele groups, offering a valuable complement to HLA typing in clinical applications.

As an application, we analyzed data from Alzheimer’s disease patients using isoLASER and report *cis*-directed splicing in multiple AD relevant genes such as *BIN1, MAPT, MSA4A6* and *HLA-DQA1*. Many of the splicing-associated variants are significantly associated with the disease based on GWAS. Disease-specific analysis unveiled differential allele-specific splicing events between AD and controls. A remarkable example is the gene *UBC,* where the linkage direction is almost completely reversed in AD compared to controls. Global alterations in splicing are increasingly appreciated in AD brains^46,58–61^, and here we present a unique approach to identify disease-specific regulation with a small cohort size. This notion calls for expanding the current focus of identifying disease-enriched mutations to disease-specific functions of genetic alterations. Lastly, around 15% of the splicing-associated variants were classified as rare or ultra-rare, according to the gnomAD database. This represents a significant expansion of the repertoire of splicing related *cis*-acting variants.

In summary, this study represents a crucial step forward in deciphering the functional impact of genetic variants on splicing using long-read RNA sequencing and the novel isoLASER method, providing valuable insights that can inform future therapeutic approaches for various human diseases.

## Methods

### Preprocessing of long-read RNA-seq data

Splice junction correction in the raw bam files was performed using TranscriptClean^62^. Initial identification of canonical splice junctions utilized GENCODE annotation. Subsequently, the function *talon_label_reads* in TALON^63^ was used to identify and remove transcripts arising from internal priming artifacts. Reads mapped to A-rich regions, defined as regions containing at least 50% adenosine bases within a 20-base window, were excluded from the analysis. Additionally, reads spanning multiple genes were also discarded to exclude potential chimeric alignments to genes with homologous regions.

Transcript structures were inferred from the sequencing reads by TALON using default parameters. Filtering steps included discarding novel transcripts supported by fewer than five reads and transcripts labeled as ‘antisense’ or ‘genomic’. Transcripts lacking splice junctions were also discarded to exclude unprocessed or short transcripts. The remaining transcripts were used for annotating individual reads and determining exonic parts. Specifically, exonic parts were defined as non-overlapping exonic segments resulting from collapsing all identified isoforms^64^. We retained only exonic parts that were not constitutively present across all transcripts, thus representing alternative splicing events. The average length of an exonic part is between 100-200 bases which is consistent with the expected length of exons in the human genome.

### isoLASER: variant calling

isoLASER’s variant caller builds on the framework used in our previous method scAllele^29^, whereby small nucleotide variants were identified using local reassembly of reads in small read segment clusters denoted as “read clusters”. We trained a multi-layer perceptron to classify variants based on multiple features, including read depth, allelic ratio, surrounding tandem repeats, and haplotype consistency. The variant quality was then calculated as the Phred-normalized probability of the False Positive classification. The performance of this classifier was evaluated using long-read RNA-seq from the GM12878 cell line and their genotyped variants from DNA sequencing. The local reassembly approach, complemented with the isoform structure determination from TALON, allows isoLASER to extract alleles and splicing information for every read enabling read-level analyses such as phasing and linkage testing.

### isoLASER: gene-level phasing

Read-level variant information extracted by isoLASER facilitates variant phasing at the transcript level while organizing reads into local haplotypes. For every expressed gene, a *v* × *n* matrix was defined where *v* was the number of heterozygous variants identified in the gene, and *n* was the number of reads aligned to the gene. Each entry contained the detected allele in the corresponding read and the variant quality score (which is the same for all reads covering the variant). A weighted k-means clustering-based approach was used to group the reads into haplotypes. This unsupervised approach used hamming distances as a measure of similarity and was further weighed by the variant quality. Best clustering was evaluated based on the average distance of the reads to the centroid. Multiple rounds with random initialization were used to mitigate the potential sensitivity of the clustering outcome to the initial conditions. Given that haplotype consistency contributes to variant quality, we performed phasing and variant scoring iteratively until convergence.

Haplotype assignment was performed within defined phasing groups, which were defined as the set of reads that overlapped at least one heterozygous variant. Typically, a phasing group encompassed all reads mapped to a specific gene. Nonetheless, a gene may contain more than one phasing group when clusters of partial reads do not share common heterozygous variants with each other.

### isoLASER: linkage test

Linkage between haplotype and local splicing at the gene level was calculated using the adjusted mutual information (AMI)^65^. The AMI of a given exonic part *i* in gene *j* is given as follows:

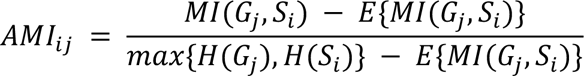

where:

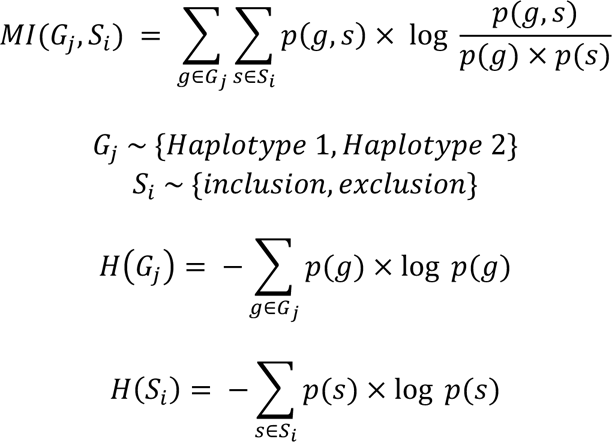

where *G_j_* represents the gene-level haplotype membership of each read, and *S_i_* corresponds to the splicing status of the exonic part *i* in each read.

The conventional mutual information approach may fail to discern meaningful linkages in cases where cluster imbalance exists, i.e., haplotype ratio of genes or inclusion ratios of exonic parts that are close to 0 or 1. AMI is less sensitive to this imbalance by subtracting the expected mutual information between two random clusterings, i.e., *E*{*MI*(*G*_*j*_, *S*_*i*_)} and normalizing by the entropy of the variables *max*{*H*(*G*_*j*_), *H*(*S*_*i*_)}.

Simulated data with randomly assigned labels (i.e., no underlying linkage) showed that AMI variance strongly depended on the total number of observations (that is, read coverage) (Fig. S1b). We modeled this variance by fitting a third-degree polynomial through the 95^th^ and 99^th^ percentile values of different coverage bins. This serves as a proxy to the AMI significance as a function of read coverage. It is also a more computationally efficient approach than bootstrapping in assigning a meaningful significance score. Based on the polynomial prediction of AMI quantiles, we categorize exonic parts into *cis*-directed if their AMI is above the 99^th^ percentile, ambiguous if they are between the 95^th^ and 99^th^ percentile, and *trans*-directed if they are below the 95^th^ percentile. Note that in analyzing actual long-read data, genes with coverage higher than 3,000 reads were down-sampled to 2,000 to reduce the processing time and memory. The observed gene coverage distribution in real data indicates most genes have significantly fewer reads than this cutoff (95% genes with ≤ 650 reads). Furthermore, as an effect size filter, we required *cis*-directed events to exhibit at least a 5% difference in PSI (delta PSI) between the two haplotypes of the same gene.

### isoLASER-joint

isoLASER can perform joint linkage analysis by merging the linkage events, and variant calls from multiple samples (Fig. S1d). For a specific exonic part, isoLASER merges the reads of the same phasing group from different samples and calculates a new AMI score. Read counts from each group are normalized to avoid confounding effects by read coverage differences.

The isoLASER-joint method affords one major advantage. When analyzing multiple samples jointly, only functional variants or variants in strong linkage disequilibrium with the functional variant will maintain a high association with the exonic parts. The association between non-functional variants and splicing is most likely inconsistent across samples. Therefore isoLASER-joint effectively “narrows down” on the candidate variants that are functional (Fig. S1d). Samples that are not testable for linkage due to low read coverage or homozygosity can also contribute to the joint analysis as they allow an examination of the consistency between alleles and splicing. Overall, as long as an alternative allele exists in the multi-sample cohort, isoLASER-joint can assess its allele-specific splicing in the specific sample and its association with splicing in the cohort, affording a possibility to study rare variants. The same AMI cutoffs as in the single-sample analysis were used to determine significance of *cis*-linkage in isoLASER-joint.

It should be noted that isoLASER (single or joint mode) has been tested on both PacBio and Oxford Nanopore long-read RNA-seq data.

### Method Benchmarking

*Variant calling:* variant calling by isoLASER was benchmarked against other Methods, such as DeepVariant, Clair3, and GATK HaplotypeCaller, following the protocol established by Souza et al^30^. In brief, precision, recall, and F1 scores were calculated at varying coverage cutoffs for all methods. Long-read RNA-seq data from WTC11 cells were used similarly as in Souza et al^30^..

*Gene-level phasing:* we compared the performance of variant phasing by HapCUT2 and isoLASER using the WTC11 data and known genome information of this cell line. Due to the potential discontinuity of the reads, both methods perform phasing in blocks. For every variant, we checked if it was phased by either method. We further assessed whether the phased variants were co-localized with other variants in the same block. We showed that blocks containing multiple variants were more easily phased by both methods. Variants phased uniquely by isoLASER tend to localize in blocks with fewer variants (Fig. 2c).

*Cis-directed event calling (linkage testing):* We compared the performance of isoLASER in identifying *cis*-directed events to that of LORALS (Github: https://github.com/LappalainenLab/lorals) which identifies allele-specific relative transcript expression. We followed the standard pipeline of LORALS with default parameters and applied it to the human tissue data from ENCODE. As this dataset lacks variant and phasing information required by LORALS, we utilized the variant calls and phasing results from isoLASER.

### Cluster assignment consistency

Hierarchical clustering (Fig. 1c, d) was performed by the package ‘aheatmap’, which uses Euclidean distance to measure the similarity between samples and the ‘complete linkage’ method for clustering. We then cut the clustering dendrogram into 7 clusters and measured their correlation with the ‘Tissue’ and ‘Donor’ labels using the variation of information (VI) metric from the R function ‘fpc::cluster.stats’. Smaller VI values represent a higher correlation between the two sets of labels.

### sQTL overlap

We downloaded sQTL results from the GTEx portal that included sGenes and significant variant-gene associations based on LeafCutter’s intron excision phenotypes. We matched similar tissues between the GTEx and ENCODE data and then overlapped the significant sQTL pairs with the splicing-associated variants and exonic parts obtained from isoLASER-joint for every tissue. Variants that were not significantly associated were included as matched controls.

There exist significant differences in how splicing is quantified by Leafcutter (used in the sQTL mapping) and isoLASER. The phenotype value used by the sQTL mapping is the junction usage within intron clusters, while isoLASER calculates PSI for each exonic part. To check the consistency between the sQTL regression parameter (beta) and isoLASER’s delta PSI value, we selected the splicing scenarios where we could directly compare these two metrics. To this end, we selected intron clusters representing exon skipping and alternative splice site usage and excluded those representing more complex splicing types, such as mutually exclusive exons or multiple exon skipping. For these scenarios, the usage of the longest junction is inversely proportional to the PSI of the skipped exon or exonic part. Thus, for Fig. 3e, we expect an inverse correlation between these two metrics.

### Gene Ontology (GO) analysis

GO terms were obtained from the Ensembl database using the R package biomaRt^66,67^. To obtain the best GO term enrichment for *cis*-directed genes, we only used genes harboring events with a delta PSI greater or equal to 0.25. The overall set of testable genes was used as background genes for each of the query gene sets. Then, for each GO term, we determined the significance of the enrichment among the query genes compared to the background using a hypergeometric test to obtain a p-value. After FDR correction using the Benjamini-Hochberg approach, only terms with FDR <= 0.05 and containing at least five query genes were considered.

### Delta delta PSI

The delta delta PSI (ddPSI) captures the difference in PSI between haplotypes and between conditions and is defined as follows:

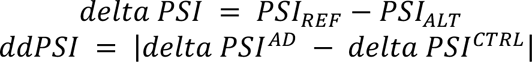

where *PSI_REF_* represents the PSI value of the exonic part linked to the reference allele, and *PSI_ALT_* represents that for the alternative allele. *ddPSI* denotes the difference in delta PSI between the AD and control samples.

The ddPSI values range from 0 to 2 representing, respectively, no difference in linkage (0) to perfect linkage in the two conditions but with the opposite alleles (2).

### HLA typing

To annotate the HLA alleles present in different samples, we mapped raw sequencing reads to the HLA allele database IPD-IMGT^68^. The alignment was performed using minimap2^69^ with the parameters ‘-ax splice:hq’ to allow for split alignment.

Supplementary and secondary alignments were excluded. We then identified the most prevalent allele groups from each HLA gene according to the standard nomenclature (https://hla.alleles.org/nomenclature/naming.html).

### AlphaFold

We followed the pipeline established in the Google Colab Notebook (https://colab.research.google.com/github/deepmind/alphafold/blob/main/notebooks/AlphaFold.ipynb) to fold the transcripts identified by TALON. This pipeline utilizes AlphaFold^70^ v.2.3.2. To assess the impact of *cis*-directed events on the resulting peptides, we specifically chose a subset of transcripts that differed solely in the inclusion or exclusion of the relevant exonic segments.

## Data availability

PacBio long-read RNA-seq of the human and mouse tissues and cell lines were obtained from the ENCODE portal (ENCODE 4 release) at https://www.encodeproject.org. In this study, we only included data generated by the PacBio Sequel II or later platforms. The Alzheimer’s ONT data were obtained from https://www.synapse.org/#!Synapse:syn52047893^45^. The GTEx sQTL data was obtained from the GTEx portal at https://www.gtexportal.org. Annotation of functional protein domains were obtained using InterPro at https://www.ebi.ac.uk/interpro.

## Code availability

The source code for isoLASER is publicly available at https://github.com/gxiaolab/isoLASER. All steps involved in detecting *cis*-directed events in this manuscript are also packaged in a WDL workflow that is publicly available at https://github.com/gxiaolab/isoLASER_paper

## Acknowledgements

We thank the Mortazavi laboratory at UC Irvine and the Wold laboratory at CalTech for producing the ENCODE dataset. We appreciate the helpful discussions with Dr. Michael R. Sawaya at UCLA. We thank members of the Xiao laboratory for helpful comments and discussions. This work was supported in part by grants from the National Institutes of Health (U01HG009417, R01AG056476, R01AG075206 to X.X.). G.Q.V. was supported by the UCLA Quantitative and Computation Biosciences Collaboratory Fellowship. K.A. was supported by the University of California-Historically Black Colleges and Universities (UC-HBCU) Fellowship. The content is solely the responsibility of the authors and does not necessarily represent the official views of the National Institutes of Health.

## Author information

### Authors and Affiliations

Department of Integrative Biology and Physiology, University of California, Los Angeles, Los Angeles, CA, USA.

Giovanni Quinones-Valdez & Xinshu Xiao

Bioinformatics Interdepartmental Program, University of California, Los Angeles, Los Angeles, CA, USA.

Kofi Amoah & Xinshu Xiao

### Contributions

G.Q.V., K.A. and X.X. conceptualized the study. G.Q.V. and K.A. performed formal bioinformatic analyses. G.Q.V. wrote the isoLASER software. X.X. provided supervision. All authors contributed to the writing of the paper and approved the final manuscript.

### Corresponding Author

Correspondence to Xinshu Xiao

